# Body size variability across habitats in the *Brachionus plicatilis* cryptic species complex

**DOI:** 10.1101/2020.04.30.070417

**Authors:** Aleksandra Walczyńska, Manuel Serra

## Abstract

The body size response to temperature is one of the most recognizable but still poorly understood ecological phenomena. Other covarying environmental factors are frequently invoked as either affecting the strength of that response or even driving this pattern. We tested the body size response in five species representing the *Brachionus plicatilis* cryptic species complex, inhabiting 10 brackish ponds with different environmental characteristics. Principal Component Analysis selected salinity and the oxygen concentration as the most important factors, while temperature and pH were less influential. Path analysis showed a positive interclonal effect of pH on body size. At the interspecific level, the size response was species and factor dependent. Under the lack of a thermos-oxygenic relationship, the expected negative response of size to temperature disappeared, but a positive response of size to oxygen remained. Our results confirm the driving role of oxygen in determining the size-to-temperature patterns observed in the field.

## Introduction

Understanding the relationship between living individuals and their abiotic environment is a fundamental step in understanding ecology. However, this scientific issue remains elusive and was quite recently even cited as one of the grand challenges of biology (Schwenk *et al.*, 2009). The most important abiotic factor in ecological niches and, consequently, species geographical ranges is temperature. It affects organisms at all levels of the organization of life, ranging from pure physical limitations (Schmidt-Nielsen, 1990; Willmer, Stone & Johnston, 2000), to ecological relationships (Begon, Townsend & Harper, 2006; Johnston & Bennett, 2008). Organisms respond to temperature in many different ways. Among these responses, the most easily recognizable and crucial for life strategies is the body size response. Size decreases with increasing temperature are both observed at the phenotypic level (Atkinson, 1994; Atkinson & Sibly, 1997) and reflected in the genetic background (Bergmann, 1847; Blanckenhorn & Demont, 2004; Blackburn, Gaston & Loder, 1999). However, this pattern is ecologically puzzling (e.g., Berrigan & Charnov, 1994), and the underlying proximate and ultimate mechanisms are not conceptually easy to test (Angilletta & Dunham, 2003). The difficulty arises from the occurrence of environmental factors correlated with temperature (Angilletta Jr, Steury & Sears, 2004), which may or may not be temperature induced. The factors that are noted in the literature as interfering with the general organismal response to temperature, which differs across habitats, are precipitation (Clusella-Trullas, Blackburn & Chown, 2011), nutrition (Horne *et al.*, 2016; Kielbasa *et al.*, 2014), seasonality (Stoks, Geerts & De Meester, 2014; Hassall, 2013), voltinism (Horne, Hirst & Atkinson, 2015), the ability to disperse (Merckx *et al.*, 2018) and oxygen availability (Rollinson & Rowe, 2018a; Santilli & Rollinson, 2018; Walczyńska & Sobczyk, 2017; Czarnoleski *et al.*, 2015). The last is of special scientific interest. A decline in oxygen availability at high temperature, especially in aquatic systems (Forster, Hirst & Atkinson, 2012), has been suggested to be a selective factor driving decreases in body size through cell shrinkage to meet aerobic metabolic demands under a diminishing oxygen supply (Woods, 1999; Verberk *et al.*, 2011). There are a number of studies confirming the size response to temperature-dependent oxygen conditions, either at the long-term genetic level or the short-term phenotypic level, both indirectly (Berner, VandenBrooks & Ward, 2007; Verberk & Atkinson, 2013; Rollinson & Rowe, 2018a; Rollinson & Rowe, 2018b; Santilli & Rollinson, 2018; Harrison, Kaiser & VandenBrooks, 2010) and directly (Czarnoleski *et al.*, 2015; Frazier, Woods & Harrison, 2001; Hoefnagel & Verberk, 2015). However, the empirical evidence confirming the adaptive significance of this pattern is very limited (Walczyńska *et al.*, 2015). Providing that the role of oxygen in the response of size to temperature is correctly predicted, the question arises of: what the actual cue for the size response is.

In this study, we examined inter- and intraspecific variability in body size in the *Brachionus plicatilis* (Rotifera) cryptic species complex in relation to environmental conditions in natural habitats. Cryptic species complexes provide a promising study system for ecological hypothesis testing because the high similarity of the species in such complexes simplifies the inference of the patterns and processes involved in evolutionary ecology. Rotifera is an especially interesting group in this regard because at least 42 species complexes of rotifers have been discovered (Gabaldon *et al.*, 2017). Among these groups, the best known is the *Brachionus plicatilis* cryptic species complex. Although 15 *Brachionus* species have been recognized using molecular techniques, only six of them have been formally described (Mills *et al.*, 2017), and four are known to inhabit ponds in eastern and central Spain. The sympatric coexistence of these species (and two additional species that have yet to be formally unnamed) in the well-documented system of brackish ponds in Spain is mediated by seasonal ecological specialization (Ortells, Gomez & Serra, 2003) related to factors such as salinity, temperature, resource use and vulnerability to predation (and Serra & Fontaneto, 2017; reviewed in Gabaldon *et al.*, 2017). A phylogenetic analysis showed signatures of coexistence in this region extending back to the Pleistocene (Gomez *et al.*, 2007). According to paleolimnological studies, this pattern has persisted for several decades, at least in two large species in the complex (*B. plicatilis sensu stricto* and *B. manjavacas*) (Montero-Pau *et al.*, 2011). According to current knowledge, between-species gene flow is absent in this complex in the wild (Gomez, Carmona & Serra, 1997). It is also important to mention that the members of the *B. plicatilis* species complex are all herbivorous and feed on different types of algae.

The first indication that *B. plicatilis* was a cryptic species complex was the observation of three apparent size classes, initially referred to as the “L” (large), “SM” (medium) and “SS” (small) morphotypes (Mills *et al.*, 2017; Serra & Fontaneto, 2017). Therefore, body size divergence and speciation are linked in the *B. plicatilis* species complex, but their causal relationship remains unknown. Regarding the response of size to temperature, a size decrease with increasing temperature has been observed at the intra- (Serrano, Serra & Miracle, 1989; Walczyńska & Serra, 2014a; Serra & Miracle, 1987) and interspecific (congeneric) levels (Gomez, Temprano & Serra, 1995; Walczyńska & Serra, 2014a). Moreover, comparison across three species from the *B. plicatilis* complex showed that species size affects the thermal dependence of diapause egg hatching (Walczyńska & Serra, 2014b). Finally, *B. plicatilis s. s.* genetically adapts to low or high temperature relatively quickly through body (and egg) size adjustment (Walczynska, Franch-Gras & Serra, 2017), revealing the crucial importance of temperature in the species’ life history, which is consistent between its phenotypic and genetic background (Walczynska, Franch-Gras & Serra, 2017).

In the present study, we exploited the potential of sediment egg banks, which were recently proposed as a newly emerging field of resurrection ecology (Weider, Jeyasingh & Frisch, 2018). In the case of *B. plicatilis*, a cyclical parthenogen, resting (diapausing) eggs are produced as a dormant, resistant stage in the bouts of sexual reproduction following periods of asexual proliferation (Serra & Fontaneto, 2017). Clones established from resting eggs deposited in the sediments of brackish ponds situated in eastern and central Spain were shown to belong to five species in the *B. plicatilis* complex differing in size. Our study aimed to reveal the role of environmental conditions in determining the body size response in the evolutionary context.

## Methods

### Limnological parameters

The system of brackish ponds in eastern and central Spain inhabited by the *Brachionus plicatilis* species complex has been sampled since the 1990s for research projects performed at the Cavanilles Institute of Biodiversity and Evolutionary Biology (ICBiBE; University of Valencia). A total of 44 ponds were seasonally inspected, and major limnological parameters (temperature, conductivity, salinity, oxygen concentration, oxygen saturation and pH) were recorded (Gomez *et al.*, 2007; Gomez, Temprano & Serra, 1995; Ortells, Gomez & Serra, 2003; Gabaldon & Carmona, 2015; Gabaldon *et al.*, 2013; Franch-Gras, Montero-Pau & Serra, 2014; Montero-Pau *et al.*, 2011; Garcia-Roger, Carmona & Serra, 2005; Lapesa *et al.*, 2002; Serra, Gomez & Carmona, 1998). Among the sampled ponds, 25 ponds were inspected at least three times in distinct periods of the year. Based on the resulting database for these ponds, Principal Component Analysis (PCA) was conducted for temperature, oxygen concentration, pH and salinity using Canoco 5.0 (Ter Braak & Šmilauer, 2012). Conductivity and oxygen saturation were excluded from the analysis because of their very high correlations with salinity and oxygen concentrations, respectively. The mean parameter values for the 25 ponds are presented in the supplementary materials (Table S1). The PCA results were used to select the ponds from which the rotifers were isolated and studied.

### Establishment of clones and body measurements

From the 25 ponds referred to above, we chose 10 ponds (Fig. 1) to study the corresponding populations of the *B. plicatilis* species complex. According to the PCA, the selected ponds represented a gradient across the two first principal components (Fig. 2). Clones of the species were established from hatched resting eggs collected from the pond sediments. We used sediments that were either previously obtained by ICBiBE or freshly collected in the field for this study. However, for two ponds, we established clones from both individuals collected in the water column and resting eggs isolated from the sediment. The details are provided together with the statistics of the pond limnological parameters in Table 1. Resting eggs were obtained from 30 g sediment samples using a modified sucrose flotation technique (Gomez & Carvalho, 2000). Hatching was induced at a salinity of 6 ppt in Petri dishes exposed to light at 25 °C. Hatchlings collected in the water column were individually transferred to the wells of a 24-well plate with 1 mL of a microalgal suspension containing approximately 3×10^5^ cells/mL of *Tetraselmis suecica* as food (under the same culture conditions as for resting egg hatching). After clonal proliferation (25 °C, 12 ppt salinity, continuous light of approximately 75 µmol quanta m^-2^ s^-1^), individuals were fixed with 40 µL of Lugol solution for size measurements. *B. plicatilis* and *B. manjavacas* were identified via the PCR-RFLP technique (Gabaldon *et al.*, 2013) in a subsample that was not fixed with Lugol. *B. ibericus* and *B. rotundiformis* were identified according to spine morphology (Ciros-Perez, Gomez & Serra, 2001). A fifth morphotype was morphologically a medium-sized *Brachionus* species (i.e., SM clade; Mills et al. 2107) in which females carried resting eggs inside their body (supplementary materials, Fig. S1). However, it was easily distinguishable from *B. ibericus*. Most likely, this morphotype was one of the not yet formally named species *B*. ‘Almenara’ or *B*. ‘Tiscar’, both of which are known to occur in the examined study area (Montero-Pau *et al.*, 2011; Gomez *et al.*, 2002). Therefore, we designated this species as SM-X. Five clones (two *B. plicatilis* or *B. manjavacas* from Hondo Sur and three *B. ibericus* or *B. rotundiformis* from Poza Norte) remained unidentified (not included in Table 1). These clones were included in the path analysis but not in the analyses in which the species factor was involved (see below).

**Table 1.** Mean values of the limnological parameters of 10 ponds from which the rotifer populations were studied, selected after PCA. Sampling details are shown. Numbers after each species name are the number of clones studied.

| Pond | Acronym | N | Temp (°C) | Salinity (g/L) | O<sub>2</sub> conc. (mg/L) | pH | Sediment collection date | Species isolated (number of clones) |
| --- | --- | --- | --- | --- | --- | --- | --- | --- |
| El Bassaret de L'Altet | ALT | 7 | 19.4 | 6.7 | 7.5 | 7.8 | 05/10/2017 | *B. ibericus* (13)*, *B. plicatilis* (6), SM-X (7)* |
| Camino de Villafranca | CVF | 5 | 20.7 | 48.8 | 10.4 | 9.2 | 22/05/2013 | *B. ibericus* (10), *B. manjavacas* (10) |
| Clot de Galvany | CdG | 5 | 18.1 | 14.1 | 9.3 | 8.3 | 05/10/2017 | *B. ibericus* (9), *B. plicatilis* (8)* |
| Hondo Norte | HON | 21 | 20.3 | 10.3 | 4.7 | 7.8 | 05/10/2017 | *B. ibericus* (6), *B. rotundiformis* (2), SM-X (9) |
| Hondo Sur | HOS | 24 | 20.6 | 10.8 | 4.4 | 7.8 | 05/10/2017 | *B. ibericus* (7), *B. manjavacas* (1), *B. plicatilis* (3), *B. rotundiformis* (2), SM-X (2) |
| Ontalafía | ONT | 3 | 14.2 | 5.4 | 9.3 | 8.6 | 22/05/2013 | *B. ibericus* (7), *B. manjavacas* (2), SM-X (5) |
| Pétrola | PET | 15 | 15.7 | 52.7 | 7.7 | 8.3 | 14/09/2017 | *B. ibericus* (10), *B. manjavacas* (8), *B. plicatilis* (1) |
| Torreblanca Poza Norte | TON | 38 | 17.8 | 10.0 | 5.0 | 7.6 | 06/06/2017 | *B. plicatilis* (10), *B. rotundiformis* (6) |
| Torreblanca Poza Sur | TOS | 10<br>6 | 18.6 | 25.3 | 10.9 | 8.0 | 06/06/2017 | *B. plicatilis* (4), *B. rotundiformis* (12) |
| Salobrejo | SAL | 12 | 14.3 | 12.9 | 5.3 | 8.9 | 24/09/2013 | *B. manjavacas* (10), *B. plicatilis* (3) |

\*sampled from the water column

**Fig. 1.**
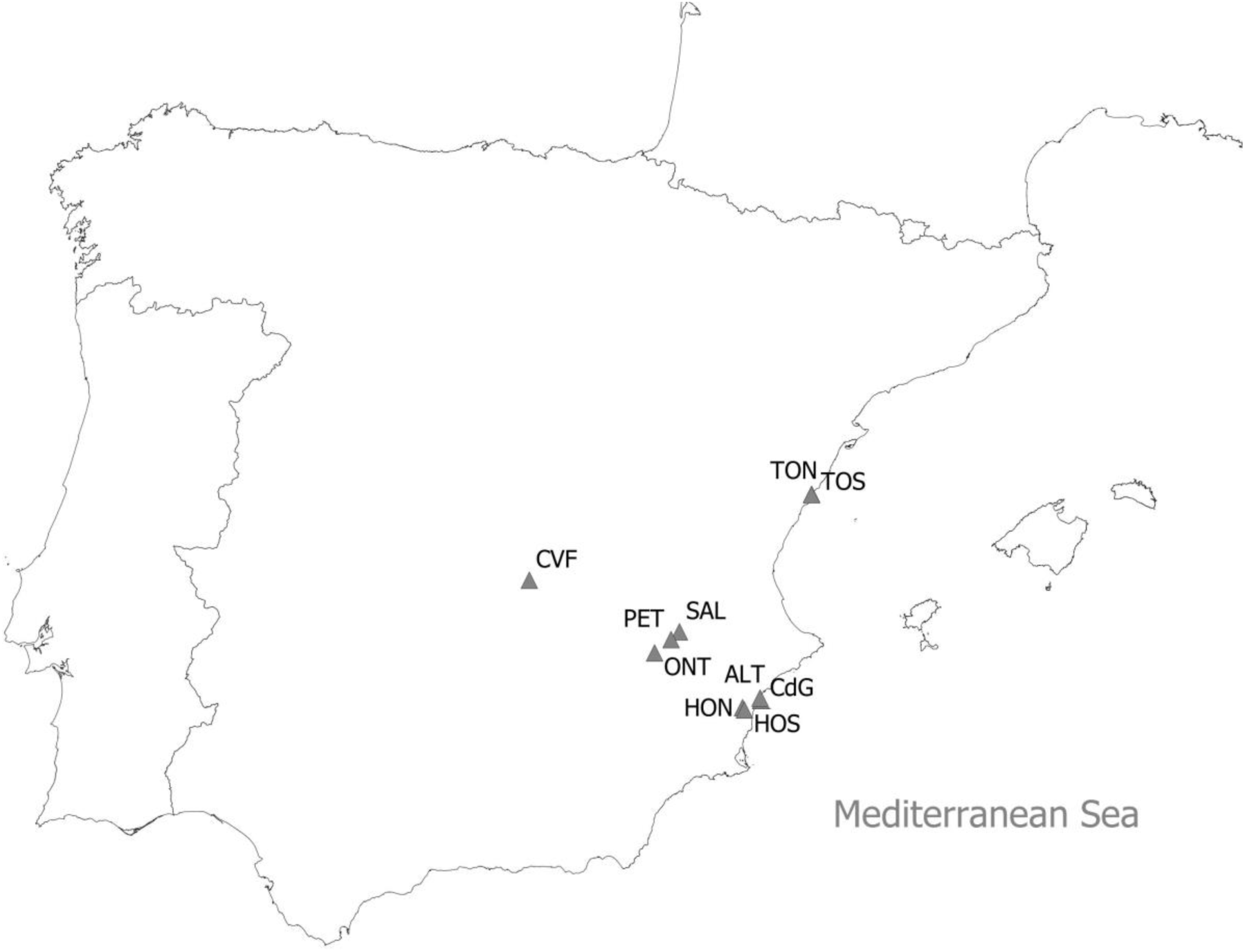
Map of Spain with the locations of the 10 ponds from which the populations of the *Brachionus plicatilis* species complex were sampled (see Table 1 for the acronyms).

**Fig. 2.**
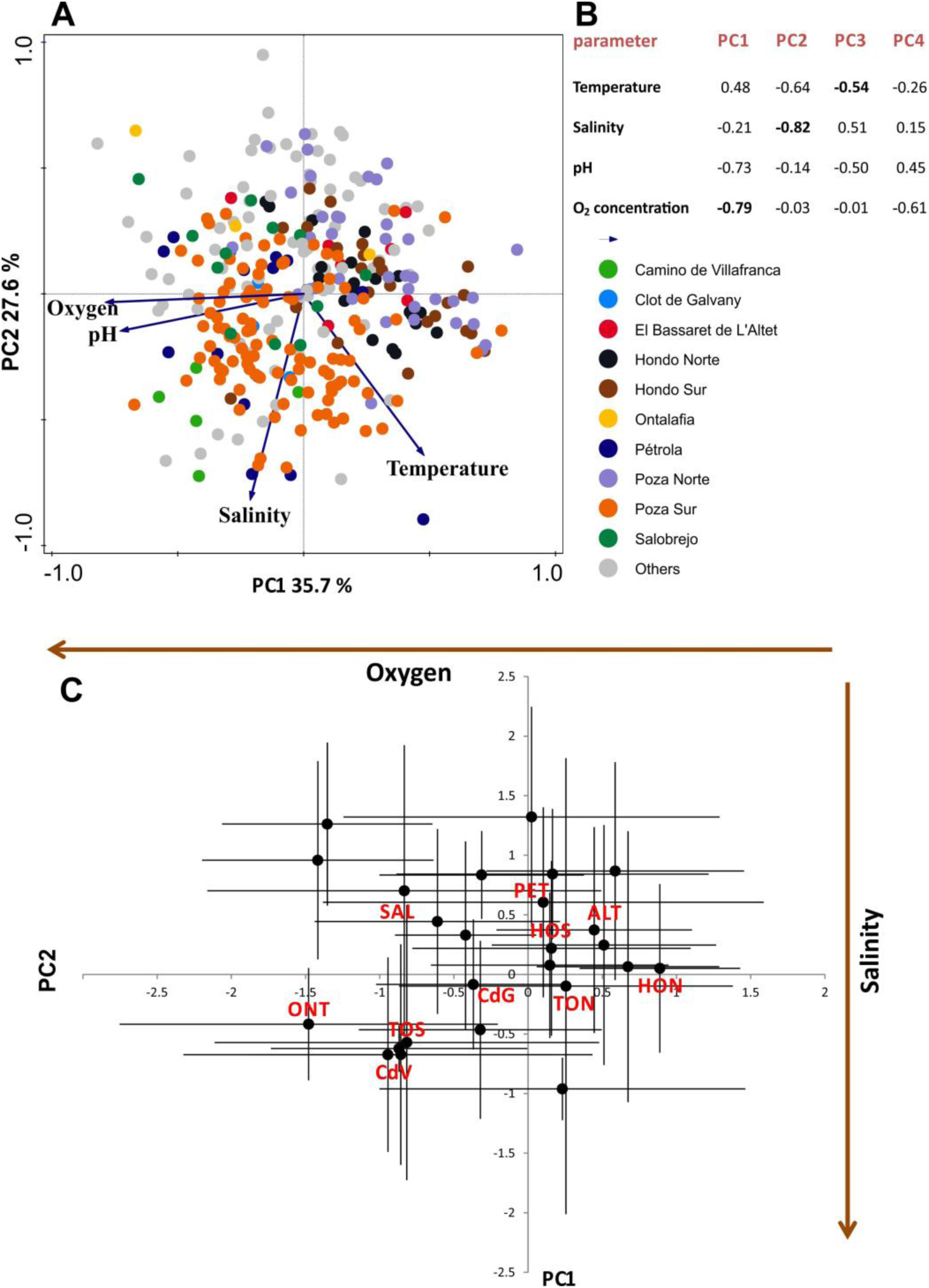
PCA of the major limnological parameters of 25 brackish ponds in eastern-central Spain where the occurrence of the *Brachionus plicatilis* species complex has been reported. **A** – PC1-PC2 scores (the 10 ponds selected to study rotifer populations are individually marked); **B** – factor loadings for PC1-PC4, where the highest load for each PC is indicated in bold; **C** – pond-specific loadings (mean ± SD) for the 10 ponds where rotifer populations were sampled (acronyms in Table 1). The arrows show the direction of the increase in the oxygen concentration (horizontal) and salinity (vertical).

For size measurements, several rotifer females from each clone carrying a single egg were photographed under 20× magnification using a Nikon Eclipse E800 microscope equipped with a Nikon DS-Ri1 camera, assisted by NIS-Elements BR software. The length and width of the lorica, the external cuticle, were measured individually in ImageJ 1.46r software (NIH, USA), and the product of these measurements served as a body size estimate; this method has been applied previously to estimate the size of *Brachionus* (Walczyńska & Serra, 2014a) and *Lecane* (Walczyńska *et al.*, 2015) rotifers. Females carrying a single egg were assumed to have matured very recently.

### Path analysis

To reveal the dependence of rotifer body size on limnological parameters, a path analysis was conducted on the measurements of a total of 178 rotifer clones, merging the species. We used PROC CALIS (SAS, 2013) with the maximum likelihood method of coefficient estimation based on the variance-covariance matrix (O’Rourke & Hatcher, 2013). First, we assumed an a priori model (referred to as the Initial model; Fig. 3A) based on common limnological knowledge (Wetzel, 2001). This model accounts for the direct effects of pH, oxygen concentrations and salinity on body size and for the direct and indirect effects of temperature.

**Fig. 3.**
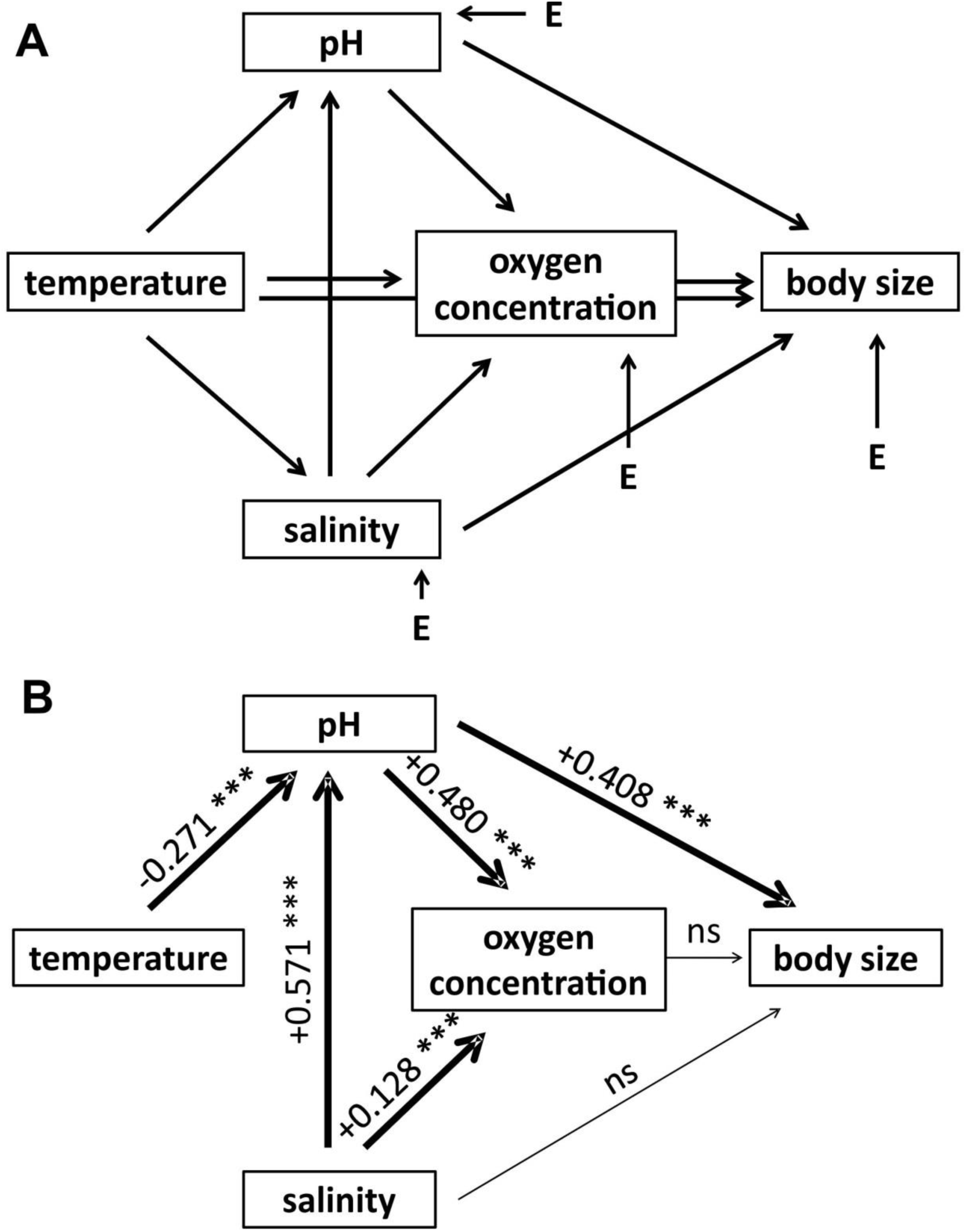
Model for the path analysis of limnological parameters affecting rotifer body size. A – the initial (a priori) model, B – path correlations with the selected model (Version 1; Table 2). Arrow thickness indicates the importance of a given path; path and covariance coefficients are provided when significant. *** p < 0.0001

Therefore, the first four factors were endogenous variables in our model, while temperature was an exogenous variable (Fig. 3A). To assess the model, we followed the rules recommended by O’Rourke and Hatcher (2013), removing the least statistically significant paths identified by the Wald test when the path-analysis model did not show proper goodness-of-fit for any of the most important indices, which are provided in Table 2A. This model selection process is a stepwise process and allowed us to achieve the reduction of the initial model and to compute the importance of each path.

**Table 2.**
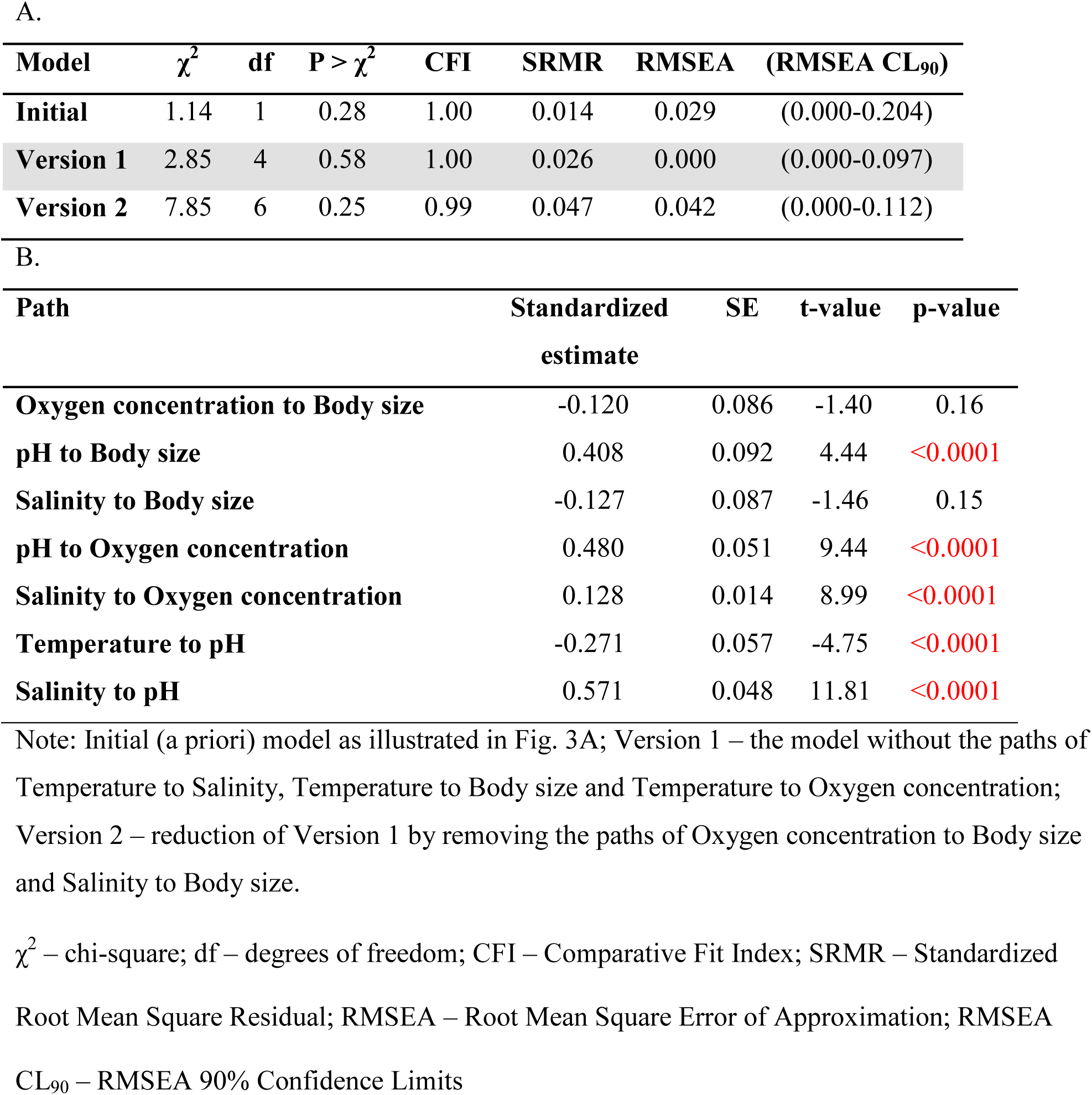
Path analysis of rotifer clone (N = 178) body size as related to the limnological parameters of 10 ponds. A – goodness-of-fit indices chosen according to recommendations of O’Rourke and Hatcher (2013); B – standardized path coefficients for the Version 1 model.

### Species-specific body size variation

The difference in body size between ponds was tested at the intra- and interspecific levels using a generalized linear mixed model (Method = REML) in PROC MIXED (SAS, 2013). The model included ‘pond’ and ‘species’ as fixed factors and ‘clone’, nested in the species and pond combination, as a random factor. Additionally, PROC REG (SAS, 2013) was used for bivariate linear regression analysis to assess the dependence of body size on temperature, the oxygen concentration, pH and salinity, addressing each species separately, with clonal mean measures as the input.

## Results

### Limnological parameters and population selection

PCA showed that 35.7% of the variance was explained by the first principal component (PC1), 27.6% by PC2, 20.0% by PC3 and 17% by PC4. The limnological parameters associated with PC1-PC3 were the oxygen concentration, salinity, and temperature, respectively (Fig. 2A, B). pH was of secondary importance in PC1 and PC3. The mean scores for the 25 ponds are presented in Fig. 2C, showing a clear transect across the first two PCs, from which the ponds whose rotifer populations were studied were selected.

### Populations and clones

In total, we measured 4150 individuals belonging to 178 clones, with 22 ± 6 individuals being measured per clone on average. The distribution of the clones across the species is provided in Table 1. The most frequent species in the studied system were *B. ibericus* and *B. plicatilis*, each of which was present in seven ponds, and the least common was *B. rotundiformis*, which was found in four ponds (Table 1). In the majority of ponds, two or three species were collected, with Hondo Sur showing the highest recorded species richness (Table 1).

### Path analysis

In the first step, the following paths were removed: Temperature to Salinity (p = 0.99), Temperature to Body size (p = 0.81), and Temperature to Oxygen concentration (p = 0.20). The output was referred to as the Version 1 model. In the next step, we removed the following paths: Oxygen to Body size (p = 0.17) and Salinity to Body size (p = 0.10), thus generating the Version 2 model. The goodness-of-fit indices were slightly worse for Version 2 than for the Version 1 model (Table 2). Therefore, we provided the final path coefficients and drew inferences for the Version 1 model (Fig. 3B, Table 2). The qualitative results did not differ between the three models. All indices obtained for the Version 1 model indicated an overall good model fit (O’Rourke & Hatcher, 2013). According to the results, (i) the body size of *Brachionus* sp. clones increased with increasing pH and was not affected by any other parameter; (ii) the oxygen concentration was positively affected by pH and salinity, with no effect of temperature; and (iii) pH was negatively affected by temperature and positively affected by salinity (Fig. 3B, Table 2B). The R^2^ value was 0.10 for body size, 0.32 for the oxygen concentration and 0.40 for pH.

### Species-specific body size variation

The mean body sizes of the species at maturity (from the smallest to the largest species) were 15 700 ± 100 for *B. rotundiformis*, 19 700 ± 100 for *B. ibericus*, 26 200 ± 200 for SM-X, 44 800 ± 400 for *B. manjavacas* and 50 800 ± 400 for *B. plicatilis* (µm^2^ ± SE). The GLM analysis showed that body size differed across the species (F_(4,146)_ = 394.44; p < 0.0001) and ponds (F_(9,146)_ = 15.84; p < 0.0001), their interaction (F_(13,146)_ = 2.96; p = 0.0007), and clones (Z-value = 7.84; p < 0.0001). The variation in body size among all five species across the studied ponds is shown in Fig. 4A, together with the pond environment described by three limnological parameters: temperature, oxygen concentration and salinity (Fig. 4B). Body size was species and parameter dependent. *B. ibericus, B. manjavacas* and *B. plicatilis* were larger in the presence of higher oxygen concentrations, and signatures of the same trend were observed for SM-X (Table 3). *B. manjavacas, B. plicatilis* and *B. rotundiformis* were larger in the presence of higher temperatures, whereas the opposite relationship was identified for SM-X. *B. ibericus, B. manjavacas* and SM-X were larger in the presence of a higher pH (Table 3). No species showed a correlation between its body size and salinity (data not shown in Table 3). The regression plots for each species-parameter combination are presented in the supplementary materials (Fig. S2).

**Table 3.** Bivariate linear regression analyses of body size on limnological parameters.

|  | *B. ibericus* |  |  | *B. manjavacas* |  |  | *B. plicatilis* |  |  | *B. rotundiformis* |  |  | SM-X |  |  |
| --- | --- | --- | --- | --- | --- | --- | --- | --- | --- | --- | --- | --- | --- | --- | --- |
|  | O<sub>2</sub> | Temp | pH | O<sub>2</sub> | Temp | pH | O<sub>2</sub> | Temp | pH | O<sub>2</sub> | Temp | pH | O<sub>2</sub> | Temp | pH |
| **slope** | 0.02 | 0.01 | 0.10 | 0.06 | 0.04 | 0.33 | 0.04 | 0.07 | -0.10 | -0.01 | 0.09 | 0.12 | 0.02 | -0.03 | 0.19 |
| **adj. R<sup>2</sup>** | 0.11 | 0.01 | 0.13 | 0.48 | 0.31 | 0.48 | 0.18 | 0.30 | 0.02 | -0.03 | 0.36 | -0.02 | 0.13 | 0.24 | 0.24 |
| **p-value** | 0.004 | 0.22 | 0.002 | <0.0001 | 0.0007 | <0.0001 | 0.006 | 0.0004 | 0.20 | 0.57 | 0.002 | 0.42 | 0.052 | 0.011 | 0.01 |

**Fig. 4.**
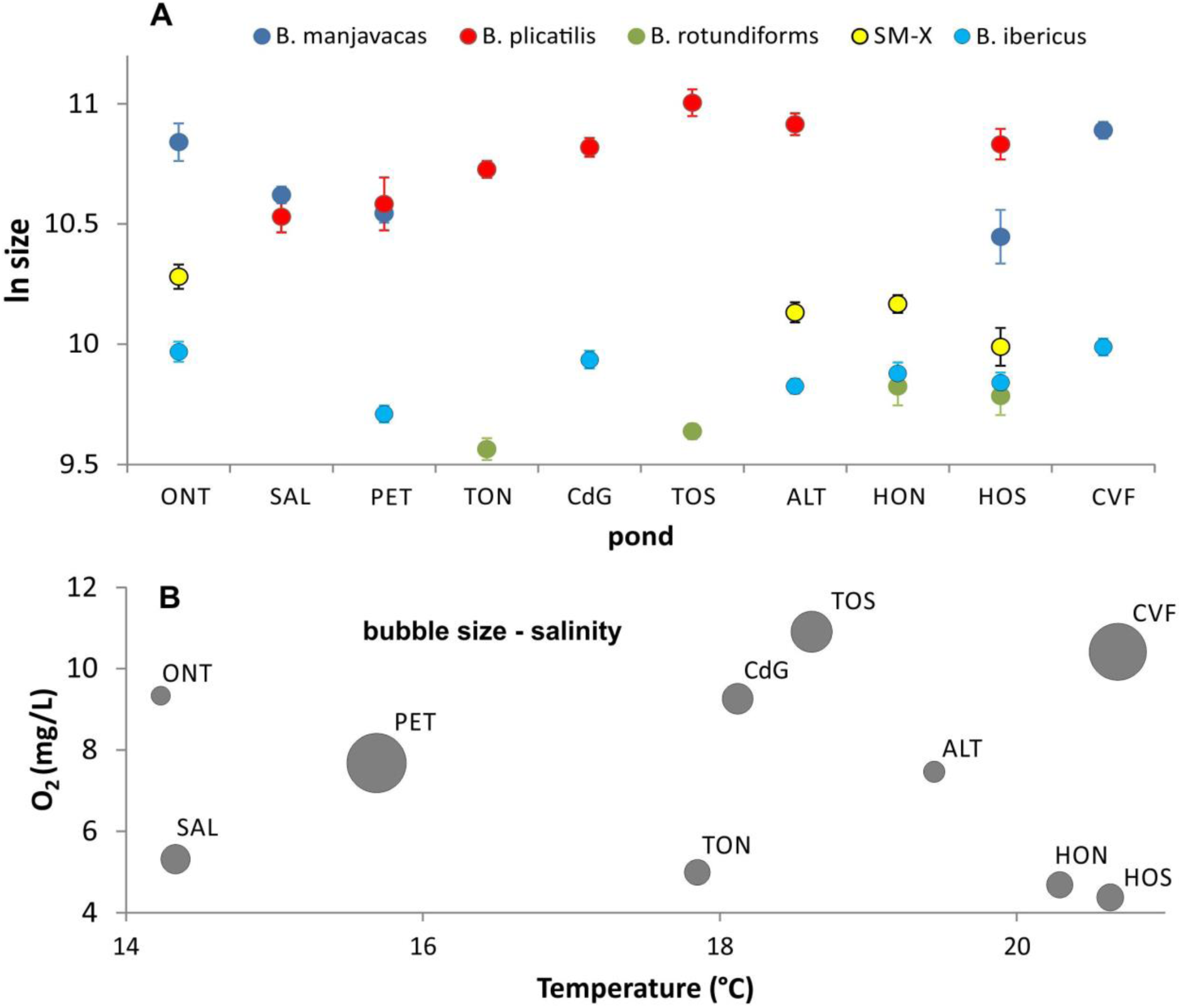
Illustration of the environmental background for body size performance across five species from the *Brachionus plicatilis* cryptic species complex from Spain. A – mean computed by averaging the clonal mean values (least square means ± SE) of the body size of five species in 10 studied ponds. B – limnological parameters of the ponds (ordered as in A) described by temperature (*x*-axis), the oxygen concentration (*y*-axis) and salinity (bubble area).

## Discussion

Based on long-term data describing the environmental characteristics of 25 shallow, brackish ponds in central and eastern Spain, we found that the most important environmental diversifying factors were oxygen and salinity, while pH and temperature were secondarily important. Five *Brachionus* rotifer species differed in body size across the ponds, with considerable variability in clones belonging to the same species. This finding suggests that the egg banks that we examined covered a wide range of variability in pond-specific environments. These results are in accordance with previous molecular marker and vital rate analyses revealing high population differentiation (Campillo *et al.*, 2011; Gomez *et al.*, 2007) and high within-population genetic diversity in species of the *B. plicatilis* complex (Gomez & Carvalho, 2000). Notably, the size variation found in our study was not due to the developmental environment, as individuals were measured under standard laboratory conditions. Size at maturity at the interclonal level (merging all the clones) was affected by the pH level, with larger clones being observed at a higher pH. Pathway analysis did not reveal a size response to the thermooxygenic conditions or salinity. A different pattern was shown when body size was analyzed separately for the studied species. In this analysis, three species exhibited larger individuals in better-oxygenated ponds, and one additional species exhibited a tendency toward this pattern (p = 0.052). Three species presented larger individuals, while one presented smaller clones in warmer ponds. Finally, three species exhibited larger individuals in the ponds with a higher pH, while none of the species showed a change in size associated with salinity. The positive response of size to the oxygen level supports the general prediction of small size being an adaptation to low oxygen availability, while the positive response to temperature contradicts the theory and the clear empirical verification of a decrease in body size with increasing temperature. Does this finding mean that the rule should be discarded? Not necessarily. The patterns that we found are likely driven by myriad abiotic and biotic factors that interact in the examined shallow, brackish water bodies, which we will discuss below.

### The effect of salinity on pH and oxygen

Salinity variation is very important in our study system, as shown by the PCA of the 25 ponds and its high relative variability across the 10 ponds in which rotifer populations were sampled (CV = 88%). Salinity affected the pH and the oxygen concentration. However, our results did not show an adjustment of the size of *Brachionus* species to salinity, confirming previous findings (Serra & Miracle, 1987), although this variable has been reported a key factor in the ecological specialization of *Brachionus* populations (Campillo *et al.*, 2011; Gabaldon *et al.*, 2015). The dependence of salinity on temperature would mean that the concentrations of all the ionic constituents dissolved in pond water would increase with desiccation caused by high temperature, as observed under summer conditions. The lack of a temperature effect on salinity led us to conclude that idiosyncratic differences in salinity between the ponds resulted from geological and geographic characteristics (e.g., proximity to the sea; (Wetzel, 2001)) and played a dominant role over seasonal changes. On the other hand, it is generally assumed that in aquatic systems, pH and oxygen concentrations are correlated with photosynthesis (Kuhl *et al.*, 1995; Wetzel, 2001). This may explain why both the PCA (the effect explained by PC1) and path analysis showed a positive relationship between pH and the oxygen concentration. Additionally, high photosynthetic activity indicates a high microalgal density, meaning that more food is available to the rotifers. This reasoning explains the positive effect of pH on the body size of *Brachionus* rotifers.

### Thermooxygenic environment

The oxygen concentration was the second most variable parameter, and temperature was the third across the investigated ponds. The overall mean oxygen concentration in the studied ponds was relatively high. Path analysis showed no effect of temperature on the oxygen concentration. This unexpected result was clarified through a bivariate regression analysis, as temperature was not negatively related to oxygen (slope = −0.06, R^2^ = 0.004, p = 0.86), contrary to the general pattern observed in aquatic systems (Wetzel, 2001; Denny, 1993). Such a negative relationship is a necessary condition for size adjustment to be responsive with regard to temperature and anticipatory with regard to oxygen (Walczyńska *et al.*, 2015). The lack of a relationship between temperature and oxygen seems to be sufficient to explain the lack of an influence of temperature or oxygen on rotifer body size that was found when individuals of the different species were merged. The importance of the link between temperature and the oxygen concentration in modulating species size was previously shown for Icelandic diatoms (Walczyńska & Sobczyk, 2017).

### Other possible influential abiotic factors

The genetic diversity of the egg banks of species of the *B. plicatilis* complex was previously found to depend on the pond area (Montero-Pau, Serra & Gomez, 2017), while the species life history strategy is affected by environmental unpredictability (Tarazona, Garcia-Roger & Carmona, 2017; Franch-Gras, Montero-Pau & Serra, 2014); specifically, interannual fluctuations in the length of the planktonic growing season (Martinez-Ruiz & Garcia-Roger, 2015). The size response to these conditions has not been systematically tested thus far.

### Sampling effects and species distribution

Our study might be affected by incomplete sampling of the species of the *B. plicatilis* complex in the 10 ponds, as additional species of the complex have been reported to occur in these ponds. For example, previous studies showed the presence of *B. ibericus* at Torreblanca Poza Sur (TOS; Garcia-Roger, 2006; Walczyńska & Serra, 2014b), while *B. rotundiformis* was previously found in El Bassaret de L’Altet (Lapesa, 2004), and the “L” morphotype (*B. plicatilis* or *B. manjavacas*) was previously known from Hondo Norte (Lapesa, 2004). On the other hand, there was no previous record of *B. rotundiformis* at Hondo Sur. The most likely reason for such differences is the habitat heterogeneity of the sediments in some ponds. Therefore, some species may be found only at specific sites within a pond. Nevertheless, our results show relatively good coverage of the general distribution of *B. plicatilis* cryptic species as reviewed by Lapesa (2004).

We show some patterns of species-specific environmental preferences. For example, SM-X is absent at high salinity, and *B. plicatilis* and *B. ibericus* are euryoic (but see below), while the occurrence of the smallest species, *B. rotundiformis*, is limited to ponds with an average temperature higher than 17 °C. This uneven species distribution may be responsible for the pattern that we found in the path analysis, showing the most apparent dependence of body size on pH, which is a proxy for resource availability. When data analyses were performed separately for each species, the two largest species, *B. plicatilis* and *B. manjavacas*, were found to share a similar strategy of exhibiting a larger body size in warmer and better-oxygenated ponds. One difference between these species was that *B. manjavacas* also presented a larger size in ponds with a higher pH. A common feature of the two SM species was positive size dependence on pH (possibly due to the direct relationship of pH with the food of these herbivorous rotifers), but *B. ibericus* was larger in higher O_2_ conditions, and its size was not related to temperature; SM-X presented a similar tendency in regard to oxygen but was smaller in warmer ponds. The smallest species (positively) responded in size solely to temperature. Such a pattern suggests that speciation within the *B. plicatilis* species complex has been driven to some extent by diverging tolerance to crucial environmental factors. It was previously found that the response of the largest *B. plicatilis* species to temperature is affected by salinity (Miracle & Serra, 1989; Serrano, Serra & Miracle, 1989) and that temperature-driven egg size adjustment occurs only at intermediate salinities (Serrano, Serra & Miracle, 1989), while unidentified SM species showed an inverse response of size to temperature regardless of salinity conditions (Serrano, Serra & Miracle, 1989). Therefore, speciation toward the “L” and “SM” morphotypes could have been associated with differential vulnerability to temperature and salinity. Our results are in line with this speculation.

### Body size and species interactions

The species in the *B. plicatilis* complex exhibit somewhat different but overlapping niches, resulting in an overlap of their seasonal distributions if they cooccur at a locality. Niche differentiation implies temperature and salinity differences (Gomez, Temprano & Serra, 1995; Gomez, Carmona & Serra, 1997), which allow seasonal succession and the partitioning of resources (Ortells, Gomez & Serra, 2003). The most interesting case is, however, the coexistence of *B. plicatilis* and *B. manjavacas*. These two largest *Brachionus* species from the complex are not distinguishable morphologically (Fontaneto *et al.*, 2007) and differ in size by only 6% (Gabaldon *et al.*, 2013). Their ecological niches are intriguingly close. Empirical evidence suggests that under environmental fluctuations involving salinity, differential adaptation to salinity and divergence in life history traits associated with different levels of opportunistic strategies are relevant to their coexistence. The results of our study indicate a new candidate as a crucial parameter affecting ecological divergence: oxygen availability. Both species adjust their size to oxygen conditions, but this relationship is steeper and stronger in *B. manjavacas*. Interestingly, this species presented the smallest body size in the least-oxygenated pond (HOS) and its largest body size in the best-oxygenated ponds (ONT and CVF). These observations may indicate high sensitivity of *B. manjavacas* to oxygen availability.

### The temperature-oxygen effect on body size

Among five *Brachionus* species collected from 10 ponds, we found only one species that exhibited a smaller body size in warmer ponds, which was consistent with the general observations regarding the relationship of size with temperature. This species, SM-X, also showed the most apparent avoidance of high salinity. The considerably variable salinity conditions could affect the response of size to temperature in the other more euryhaline species. However, the most likely factor responsible for the reversal of the response of size to temperature in our study system is the lack of the common, expected negative temperature-oxygen relationship. The increase in body size with increased nutrition, which was the most apparent result of our path analysis, could mask the possible pattern of decreasing size with increasing temperature, rather than explaining the generally reversed pattern that we found. As noted elsewhere, “*the stronger effect of nutrition than of oxygen may be observed when temperature-dependence of the former is steeper than that of the latter*” (Walczyńska & Sobczyk, 2017). Therefore, in our study system, salinity (indirectly) and pH (directly) affected the response of size to temperature, causing the absence of the expected decrease with increasing temperature. Intriguingly, under such conditions, the rotifer species exhibited smaller sizes at lower oxygen levels, as predicted by the theory, confirming the crucial role of the oxygen concentration in driving body size patterns (Walczyńska & Sobczyk, 2017). The only exception to the “when there is less oxygen, grow smaller” rule was observed for the smallest species, *B. rotundiformis*. However, as mentioned above, this species occurred only in warm waters, and possibly because of its small size, it is equipped with physiological mechanisms for dealing with hypoxia, making the body size plasticity found in other species unnecessary. The relationship of the strength of the response of size to temperature with the level of species thermal specialization was noted in another study on three *Brachionus* species, in which only *B. plicatilis*, the largest species, which is euryoic, showed a phenotypic size decrease with increasing temperature, while two other less-temperature-tolerant species, *B. ibericus* and *B. rotundiformis*, showed no such pattern. Although we found *B. ibericus* to be the most frequently occurring species across ponds, this species shows thermal specialization by occurring in the water column within the narrowest time window during the year (Gomez, Temprano & Serra, 1995).

Previous studies confirm that our results do not violate the general size-to-temperature rules for *Brachionus* rotifers. Clear size-dependent temperature preferences were shown in three *Brachionus* species originating from the same pond system: (i) smaller species generally presented a higher optimal temperature in relation to the population growth rate (Walczyńska & Serra, 2014a); (ii) smaller species preferred higher temperatures for hatching from resting eggs (Walczyńska & Serra, 2014b); and (iii) *Brachionus plicatilis s. s.* decreased in size with increasing temperature, which was reflected at the levels of both short-term phenotypic, nongenetic plasticity (Walczyńska & Serra, 2014a; Serra & Miracle, 1987) and genetics (Walczynska, Franch-Gras & Serra, 2017).

## Acknowledgments

We are grateful to Eduardo García-Roger and María José Carmona for the identification of some *B. rotundiformis* and *B. ibericus* clones, to Ana Hidalgo for her constant help and advice throughout the lab work, to Łukasz Sobczyk for conducting the PCA and to Aleksandra Pępkowska-Król for creating a map with the pond locations. The work was supported by the National Science Center of Poland (OPUS 2015/19/B/NZ8/01948) and by Jagiellonian University (DS/INoS/757/2019). The authors declare no conflict of interest.

## Notes

### Competing Interest Statement

The authors have declared no competing interest.

