## Supplemental Figures and Tables for "Body size variability across habitats in the *Brachionus plicatilis* cryptic species complex"

**Table S1.** The mean values for the limnological parameters for 25 ponds in central and eastern Spain inhabited by the *Brachionus plicatilis* cryptic species complex that were initially examined in this study. Ponds in which the sediment egg banks were examined are bolded. N – the number of records per pond.

| pond | Temp (°C) | Salinity (g/L) | pH | $O_2$ (mg/L) | N |
|---|---|---|---|---|---|
| Almenara | 20.5 | 2.5 | 8.1 | 8.2 | 15 |
| **Camino de Villafranca** | **20.7** | **48.8** | **9.2** | **10.4** | **5** |
| Canal Central | 13.6 | 3.2 | 7.8 | 7.8 | 12 |
| Chiprana | 17.8 | 37.8 | 8.8 | 11.6 | 9 |
| **Clot de Galvany** | **18.1** | **14.1** | **8.3** | **9.3** | **5** |
| **El Bassaret de L'Altet** | **19.4** | **6.7** | **7.8** | **7.5** | **7** |
| Estany de Cullera | 19.5 | 18.1 | 8.2 | 6.9 | 3 |
| Gallocanta | 15.6 | 117.0 | 8.6 | 6.5 | 3 |
| Grande de Villafranca | 19.4 | 6.9 | 8.5 | 9.6 | 7 |
| **Hondo Norte** | **20.3** | **10.3** | **7.8** | **4.7** | **21** |
| **Hondo Sur** | **20.6** | **10.8** | **7.8** | **4.4** | **24** |
| Horna | 17.4 | 3.4 | 8.7 | 10.2 | 3 |
| Hoya del Monte | 19.6 | 14.2 | 8.6 | 7.4 | 4 |
| Hoya Elvira | 15.8 | 4.4 | 7.9 | 5.1 | 3 |
| Hoya Rasa | 15.9 | 36.8 | 9.1 | 5.3 | 3 |
| Hoya Yerba | 17.0 | 8.1 | 8.1 | 7.3 | 3 |
| La Campana | 13.9 | 5.4 | 9.3 | 10.7 | 3 |
| Manjavacas | 13.8 | 22.7 | 9.1 | 10.6 | 4 |
| **Ontalafia** | **14.2** | **5.4** | **8.6** | **9.3** | **3** |
| P. Muñoz | 12.2 | 4.4 | 8.8 | 5.1 | 4 |
| **Pétrola** | **15.7** | **52.7** | **8.3** | **7.7** | **15** |
| **Poza Norte** | **17.8** | **10.0** | **7.6** | **5.0** | **38** |
| **Poza Sur** | **18.6** | **25.3** | **8.0** | **11.0** | **106** |
| **Salobrejo** | **14.3** | **12.9** | **8.9** | **5.3** | **12** |
| Taray | 19.4 | 4.1 | 8.4 | 10.1 | 5 |

**Fig. S1.** An individual of the *Brachionus* sp. designated SM-X in our study. A female is carrying a resting egg internally, a phenomenon previously reported for another SM (medium-sized) species, *Brachionus ibericus*. Clone established from El Bassaret de L'Altet pond.

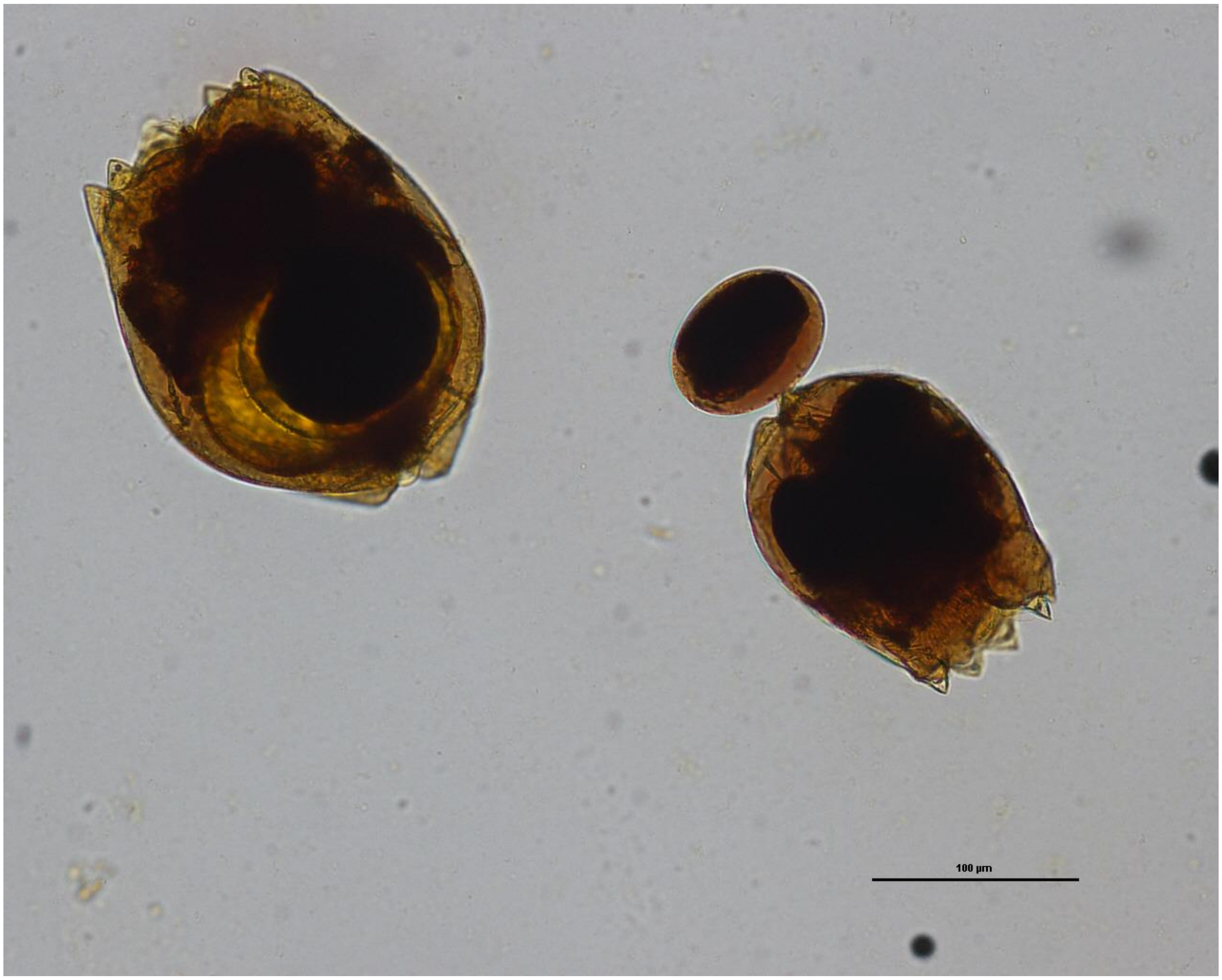

**Fig. S2.** Simple regressions of species body size (in ln) and four environmental factors: the oxygen concentration, salinity and temperature (driving the environmental PCA) and pH, which was of minor importance in that analysis but affected the body size of some species. The fitting line with the 0.95 Confidence Intervals shown only for significant relationships.

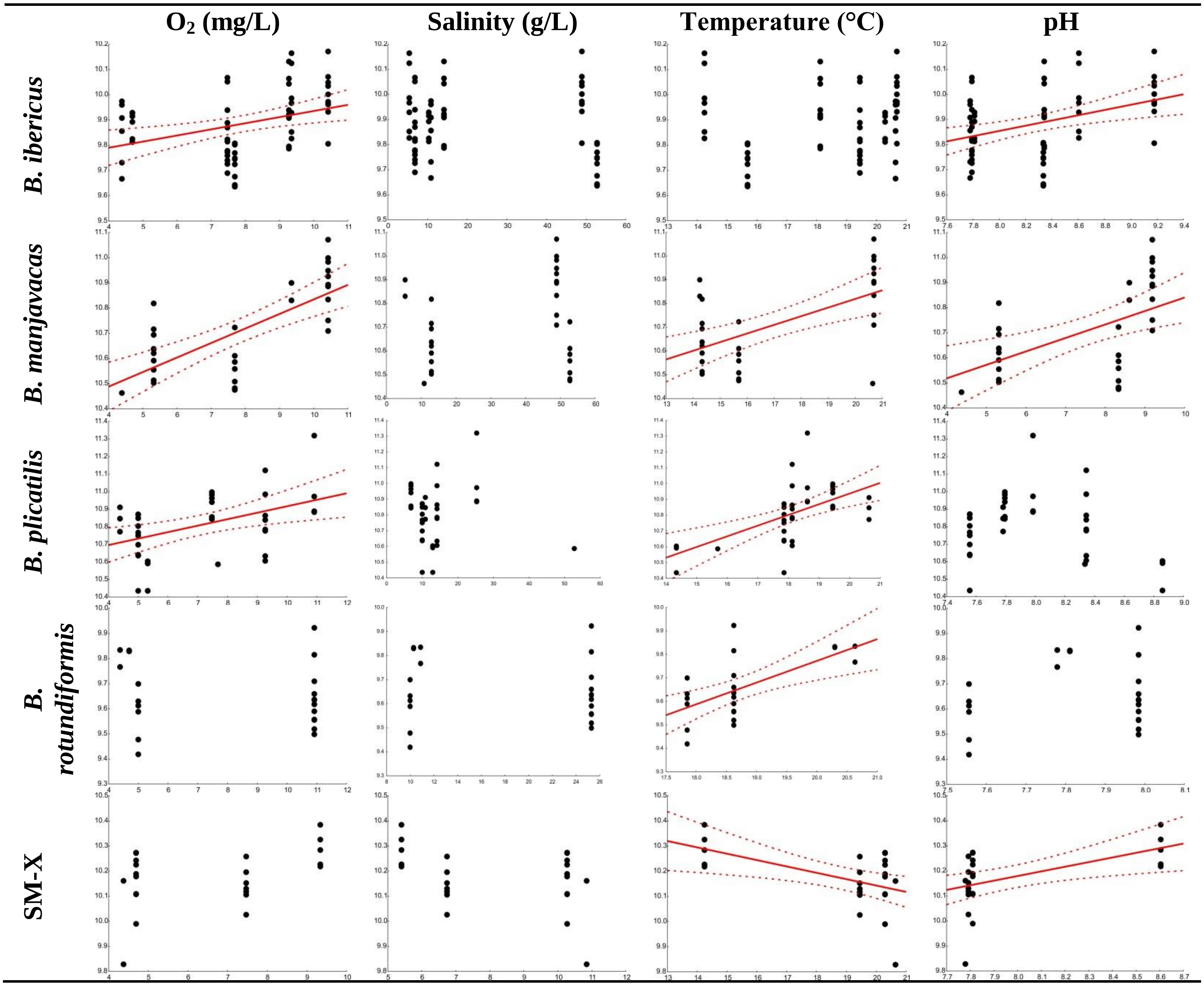
